## Supplementary figures 1 to 11, Supplementary tables 1 and 2 for "Methionine aminopeptidase 2 and its autoproteolysis product have different binding sites on the ribosome"

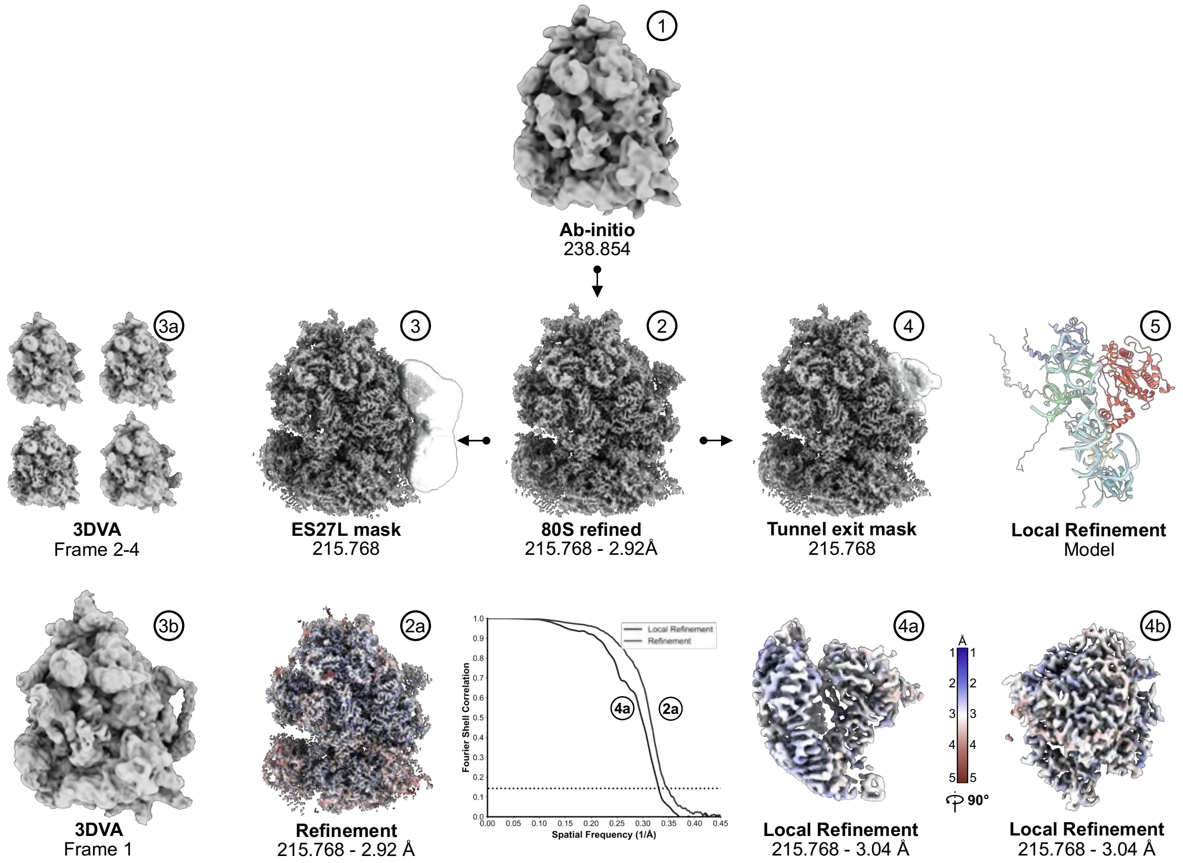

**Supplementary Figure 1:** Cryo-EM data processing of *Hs*MAP2 on 80S ribosomes using CryoSPARC. After pre-processing and three rounds of 2D classification, selected particles were subjected to Ab-initio reconstruction **(1)** to obtain a preliminary 3D volume. After further classification using Heterogeneous refinement, residual particles were used to perform Homogenous refinement, yielding a refined map of 2.92 Å resolution **(2)**. Masks were placed around the tunnel exit and ES27L **(3)** to perform focused 3D variability analysis. The output of this job was passed through a 3D variability display job into 5 particle clusters. All clusters were individually subjected to Ab-initio reconstruction, followed by Homogenous refinement. In one of the particle subset, the refined map showed the position of ES27L **(3b)**, while the others lacked this feature **(3a)**. Using a mask around MAP2, a local refinement was performed **(4)**, yielding a local resolution of 3.04 Å **(4a&b)**. Local resolution estimations were performed on the overall refinement **(2a)**, as well as the local refinement **(4a)**. Additionally, FSC curves were plotted for these two refinement. Both refinements were used to build a model of MAP2 at the ribosomal tunnel exit **(5)**.

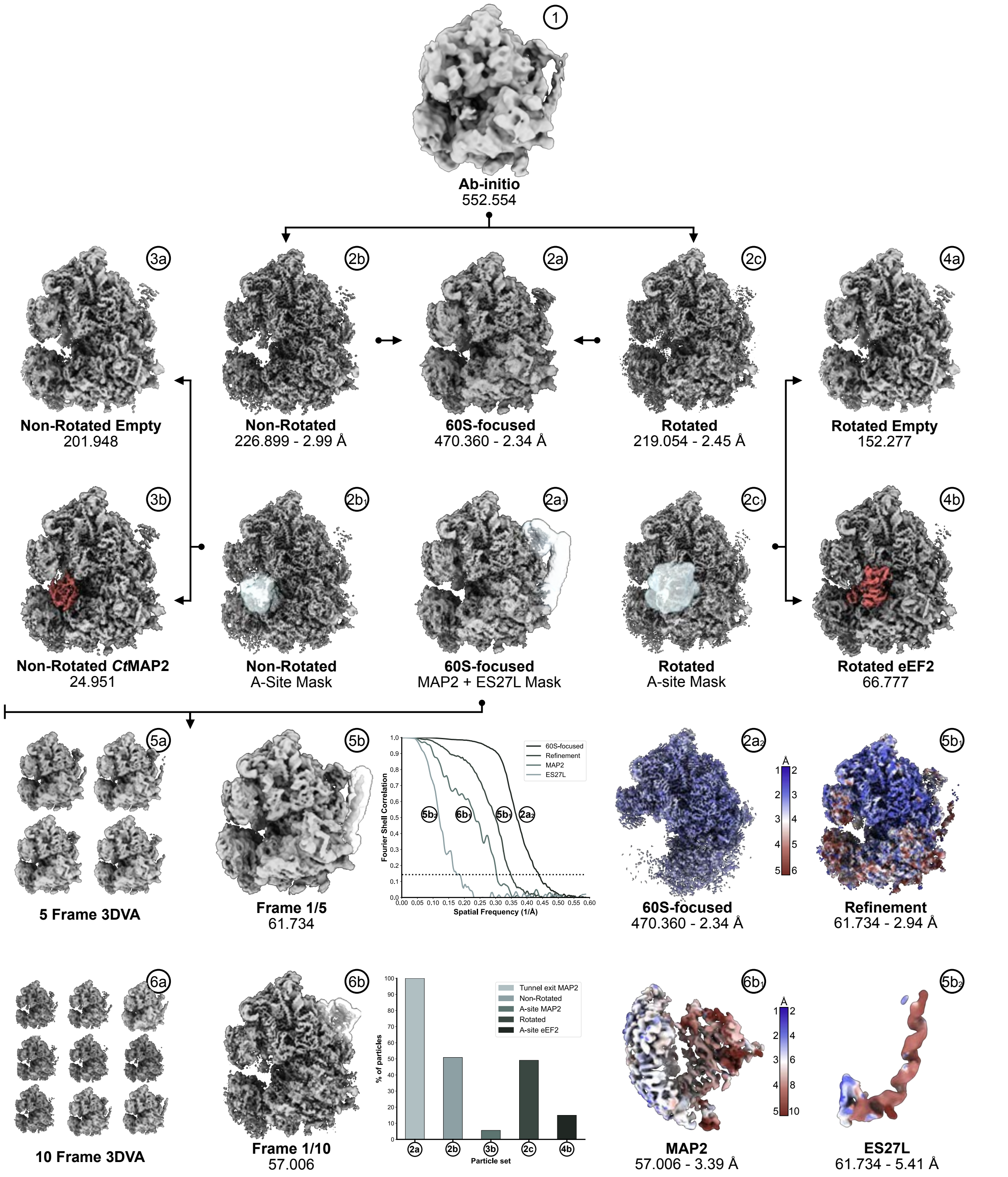

**Supplementary Figure 2:** Cryo-EM data processing of *Ct*MAP2 on the ribosome using CryoSPARC. After pre-processing and 3 rounds of 2D classification, selected particles were subjected to Ab-initio reconstruction **(1)** to obtain a preliminary 3D volume. Heterogeneous refinement revealed that the 40S subunit was rotated in nearly half of all 80S particles. Particles in the rotated- and non-rotated state were individually subjected to Homogenous refinement to obtain resolutions of 2.45 Å **(2c)** and 2.99 Å **(2b)** respectively. A mask was placed in the A-site **(2b1, 2c1)**, to perform local classification using 3D variability analysis. Particles were subsequently output into two clusters, along variability component 1, and subjected to Ab-initio reconstruction and Homogenous refinement. In ~11% (24951 particles) of all particles in the non-rotated state, MAP2 could be identified in the A-site **(3b)**, while the A-site was unoccupied for the remaining particles **(3a)**. The same procedure was performed for the A-site of 80S particles in the rotated state. Here, ~30% of ribosomes carried eEF2 **(4b)**, while the A-site was unoccupied for the remaining particles **(4a)**. Since the rotation of the 40S subunit had no apparent effect on the MAP2 interaction at the tunnel exit, particles from both subsets were merged and subjected to another Homogenous refinement **(2a)** reaching a resolution of 2.34 Å at the 60S subunit **(2a)**. Focused 3D variability analysis was performed with a mask around ES27L and MAP2, and particles were output into 5 and 10 clusters. Each particle cluster was subjected to Ab-initio reconstruction and Homogenous refinement. In cluster 1/5 **(5b)** ES27L was best define, and a mask was placed around ES27L for a subsequent local refinement **(5b2)**. Likewise, cluster 1/10 **(6b)** revealed the highest resolution MAP2 density. A mask was placed around MAP2 and a local refinement was performed **(6b1)**. Local resolution estimations were performed for all maps that were used for model building **(2a1, 5b1, 5b2, 6b1)**, and FSC curves are plotted. The final model was refined into map 5b1. The abundance of particles in the rotated and non-rotated state, as well as the occupancy of the A-site and tunnel exit are shown in a bar plot.

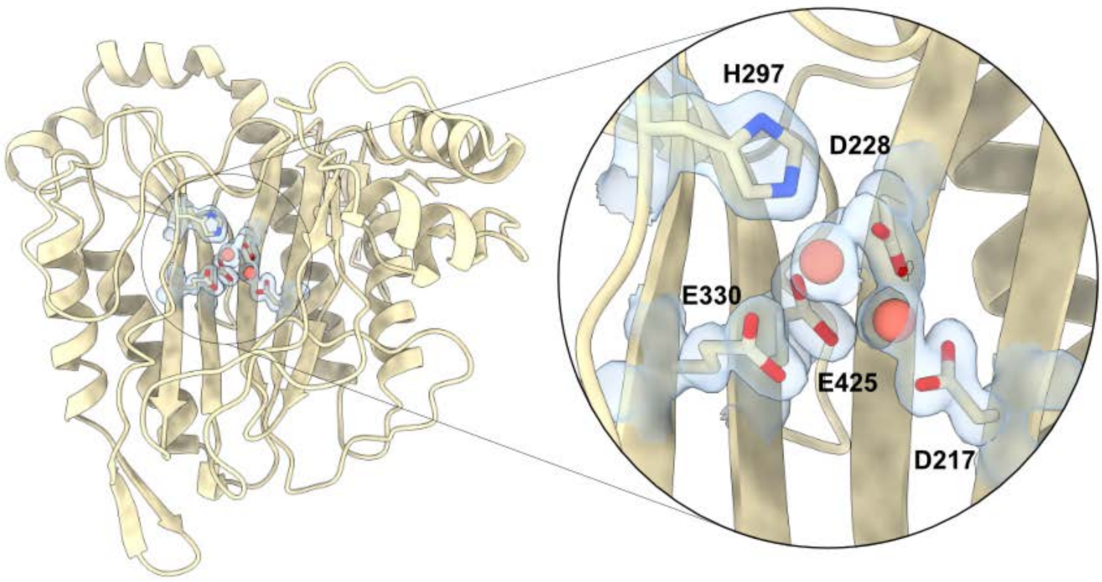

**Supplementary Figure 3:** Crystal structure of *Ct*MAP2ΔN at 1.3 Å resolution. The inset shows the active site with the five conserved catalytic residues. Two metal co-factors are coordinated by these residues. The density is shown around the side chains, and the metal cations.

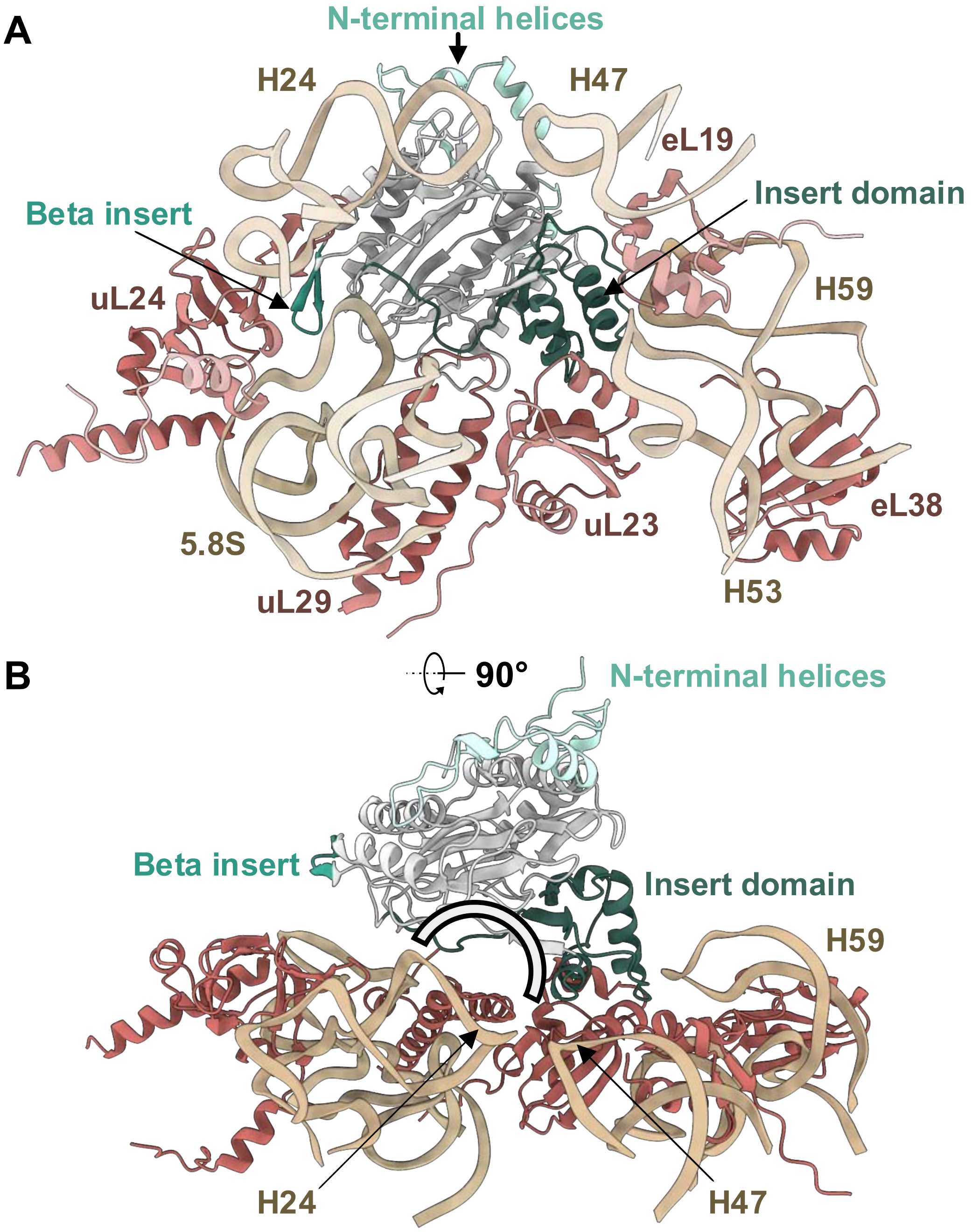

**Supplementary Figure 4:** Binding mode of MAP2 on the ribosomal tunnel exit. **(A)** View from inside the tunnel exit into the active site of *Ct*MAP2. The three main features and insertions that distinguish MAP2 from the plain pita bread fold of MAP1 are highlighted in green. The Insert domain forms the main contact at the tunnel exit, while the Beta insert and N-terminal helices do not contribute to tunnel interaction. **(B)** Side view on the MAP2-80S interaction. The MAP2 vestibule sits centrally on the tunnel exit and is emphasized by half circle.

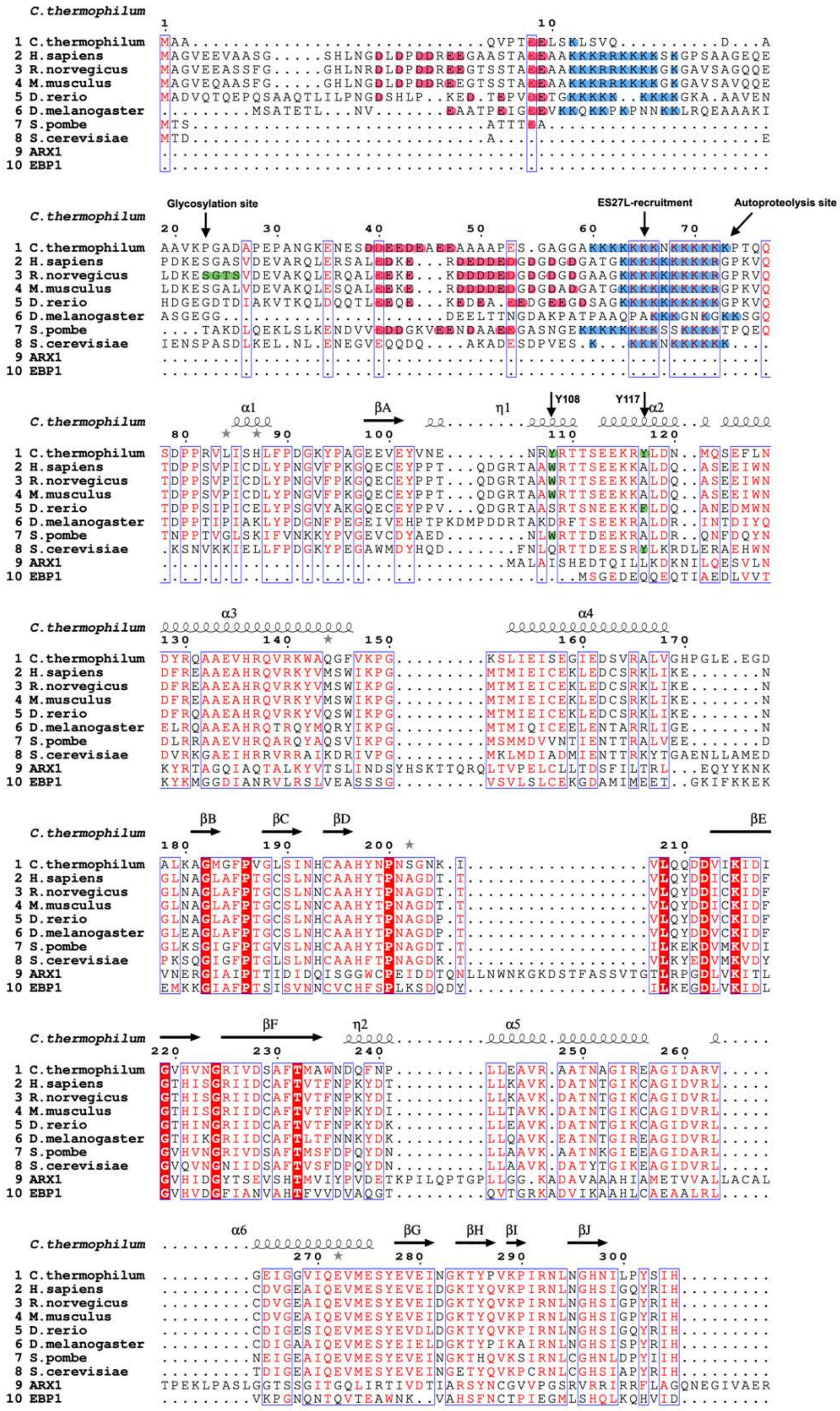

**Supplementary Figure 5A:** Multiple sequence alignment of MAP2 from 8 different organisms, as well as MAP2 like proteins *Sc*Arx1 and *Hs*Ebp1. Secondary structure elements are shown for *Ct*MAP2 above the sequences, as present in the crystal structure. Conserved residues are highlighted in red. Poly-ionic regions at the N-terminus of MAP2 are highlighted in blue (cationic) and pink (anionic). The cationic region required for ES27L recruitment is indicated by an arrow, as is the predicted autoproteolysis site. The glycosylation site at the N-terminus of RnMAP2 is highlighted in green. Important residues for A-site binding of *Ct*MAP2 (Y108 and Y117) are also indicated by an arrow.

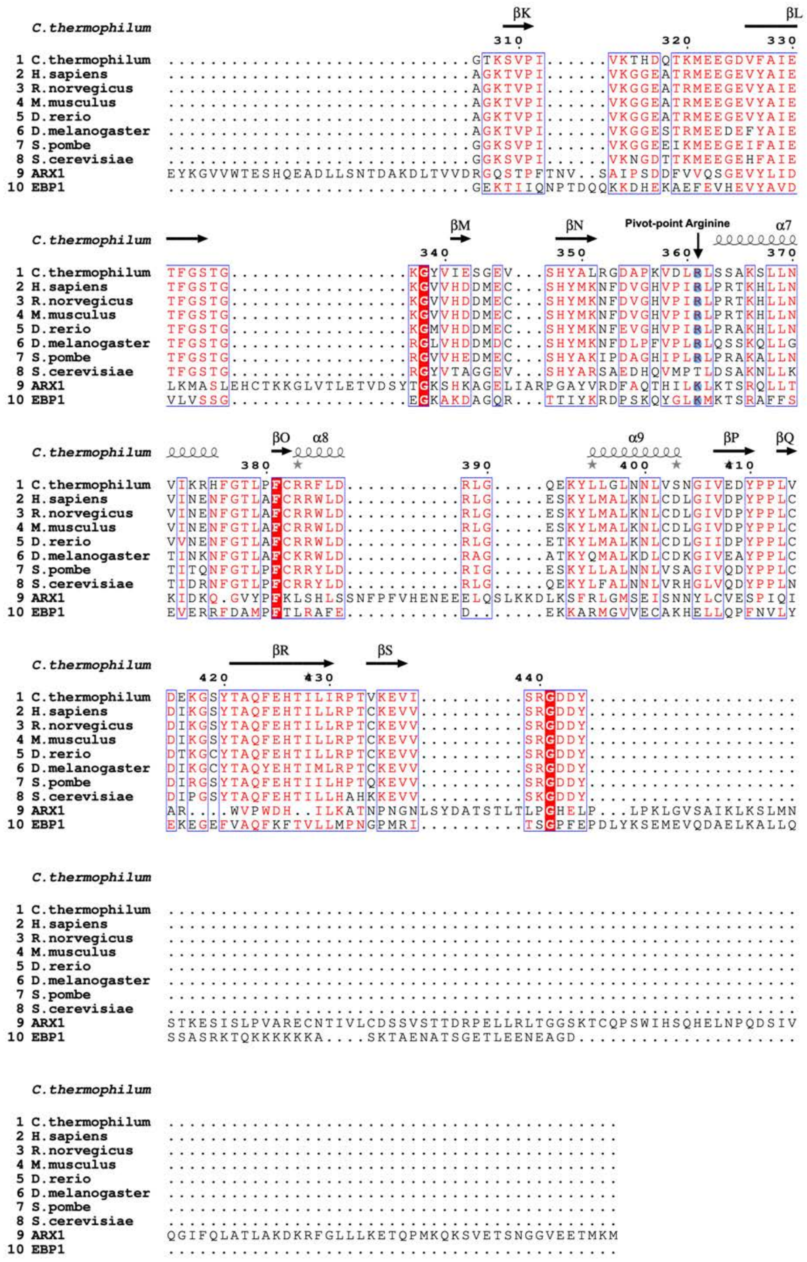

**Supplementary Figure 5B:** Multiple sequence alignment of MAP2 from eight different organisms, as well as MAP2 like proteins *Sc*Arx1 and *Hs*Ebp1. Secondary structure elements are shown for *Ct*MAP2 above the sequences, as present in the crystal structure. Conserved residues are highlighted in red. The conserved arginine that serves as the pivot point for MAP2 rotation is highlighted in blue.

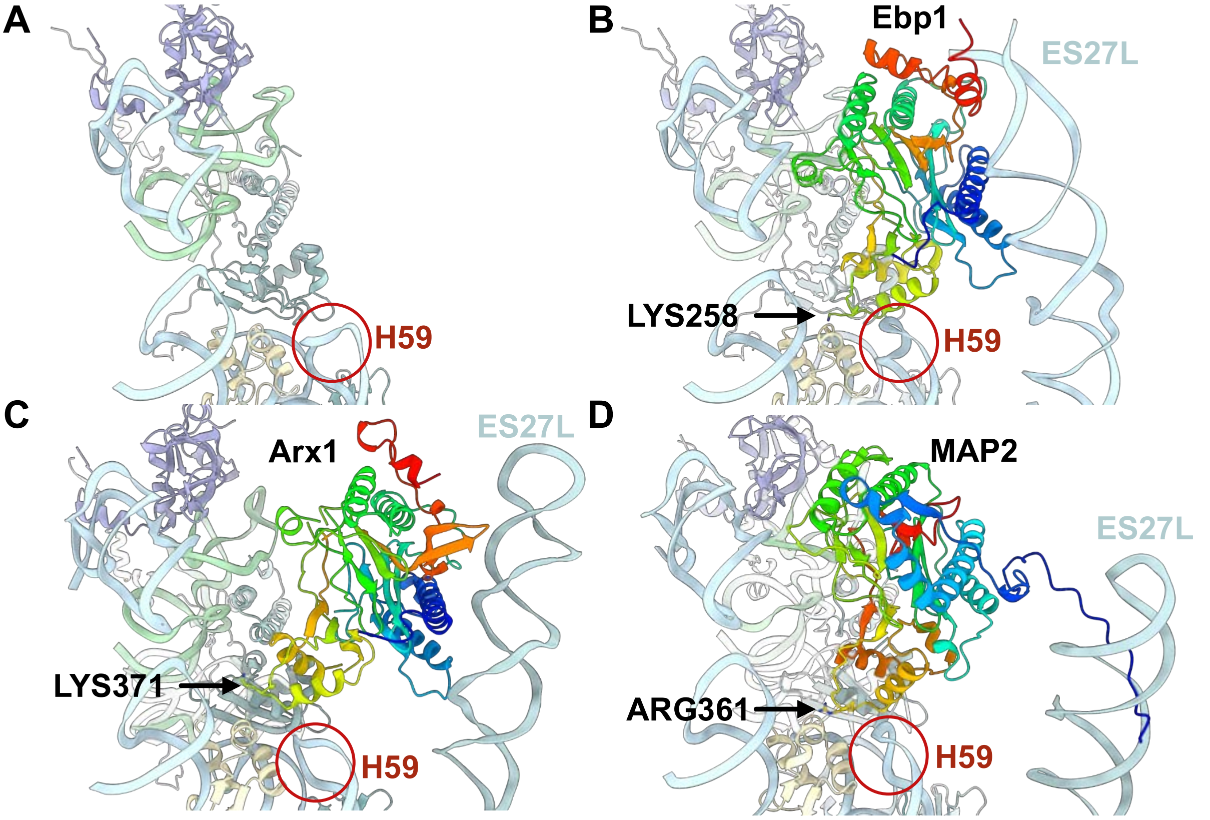

**Supplementary Figure 6:** Comparison of the binding mode between *Hs*Ebp1, *Sc*Arx1 and *Ct*MAP2. The proteins of interest are colored from blue (N-terminus) to red (C-terminus) **(A)** In the absence of a MAP2-like protein, the structure of H59 from H. sapiens ribosomes is shown. **(B)** The recruitment of HsEbp1 induces a remodeling of H59. Unlike MAP2 and Arx1,Ebp1 has two contact sites to ES27L, at the C-terminus and the N-terminus. The Ebp1-ES27L interaction is more rigid and robust than the MAP2-ES27L. The conserved pivot point arginine that is found in catalytically active MAP2s is not present in Ebp1, instead it has a lysine in this position (LYS258). **(C)** ScArx1 also recruits ES27L with its N-terminus, similar to Ebp1, but only shows one contact to the expansion segment. The pivot point arginine found in MAP2 is also missing (LYS371 instead). **(D)** MAP2 binding is highly dynamic and also recruits ES27L. Unlike Ebp1, it only binds ES27L with the unstructured N-terminal extension. CtMAP2 does not remodel H59, as observed for HsMAP2 and HsEbp1. Arg361 acts as the point of rotation at the tunnel exit.

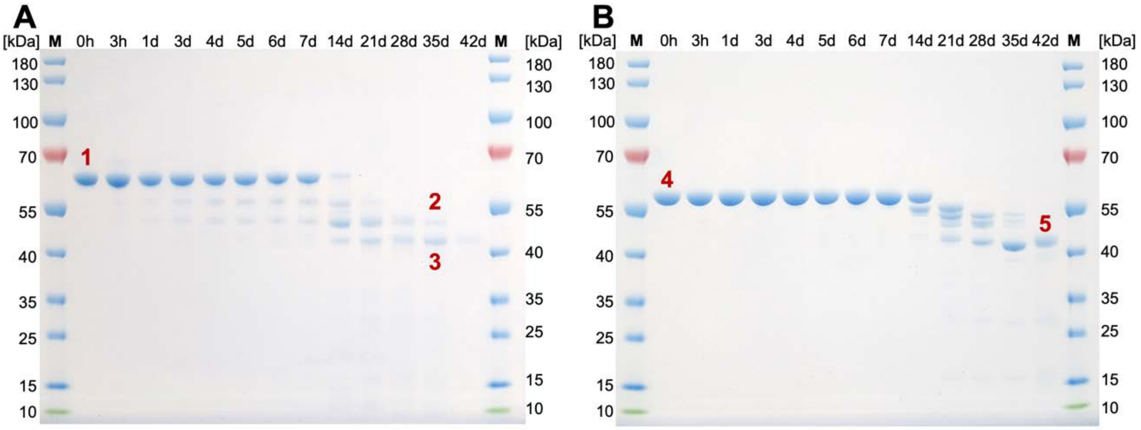

**Supplementary Figure 7:** Degradation of *Hs*MAP2 **(A)** and *Ct*MAP2 **(B)** at 4°C. MAP2 was purified from insect cells and stored at 4°C at a concentration of 1 mg/ml. Samples were collected at different intervals and analyzed by SDS-PAGE. **(A)** Full length *Hs*MAP2 (band 1) starts degrading after 3 days and yields two bands after 35 days (bands 2 & 3). Band 3 corresponds to the core domain at 41.5 kDa. **(B)** *Ct*MAP2 shows enhanced stability and only starts degrading after 14 days. *Ct*MAP2 degrades into a single stable band at 41.3 kDa, corresponding the core domain.

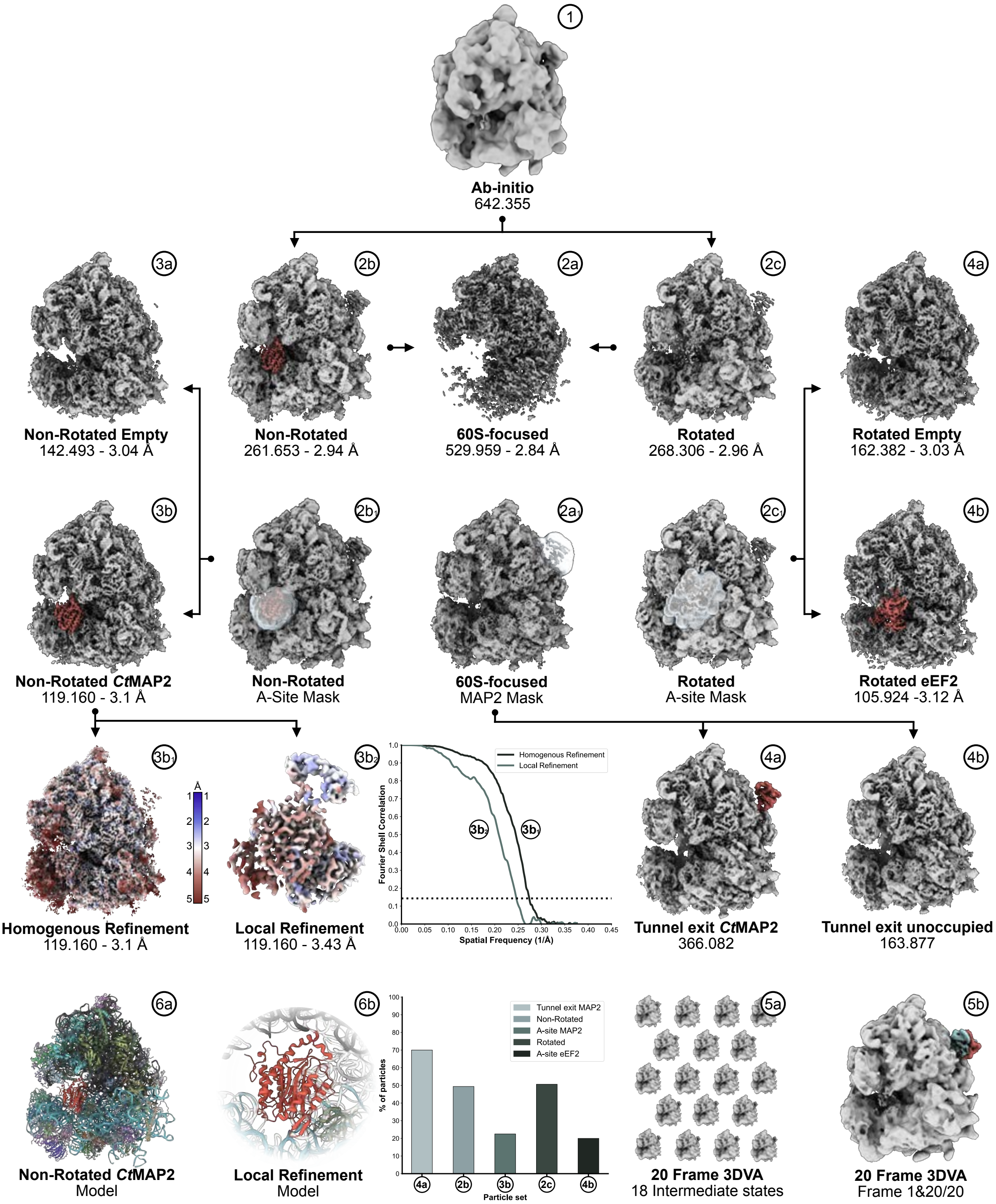

**Supplementary Figure 8:** Cryo-EM data processing of *Ct*MAP2ΔN on the ribosome using CryoSPARC. After pre-processing and 3 rounds of 2D classification, selected particles were subjected to Ab-initio reconstruction **(1)** to obtain a preliminary 3D volume. Heterogeneous refinement revealed that the 40S subunit was rotated in nearly half of all 80S particles. Particles in the rotated- and non-rotated state were individually subjected to Homogenous refinement to obtain resolutions of 2.96 Å **(2c)** and 2.94 Å **(2b)** respectively. A mask was placed in the A-site **(2b1, 2c1)**, to perform local classification using 3D variability analysis. Particles were subsequently output into two clusters, along variability component 1, and subjected to Ab-initio reconstruction and Homogenous refinement. In ~46% (119160 particles) of all particles in the non-rotated state, MAP2 could be identified in the A-site **(3b)**, while the A-site was unoccupied for the remaining particles **(3a)**. The same procedure was performed for the A-site of 80S particles in the rotated state. Here, ~39% of ribosomes carried eEF2 **(4b)**, while the A-site was unoccupied for the remaining particles **(4a)**. Since the rotation of the 40S subunit had no apparent effect on the MAP2ΔN interaction at the tunnel exit, particles in the rotated and non-rotated state were merged and subjected to another Homogenous refinement **(2a)** reaching a resolution of 2.84 Å at the 60S subunit **(2a)**. Focused 3D variability analysis was performed with a mask around MAP2ΔN, and particles were output into 2 and 20 clusters. When outputting particles into 2 clusters along variability component 1, particles that had MAP2ΔN on the tunnel exit **(4a)** could be separated from those where the tunnel exit was unoccupied **(4b)**. 3D variability analysis into 20 clusters (5a+b) revealed that MAP2ΔN undergoes a rotation at the tunnel exit **(5b)**. Using the particles that had MAP2 in the A-site **(3b)**, a local refinement was performed. Subsequently, Local resolution estimations were performed for the overall refinement and the local refinement **(3b1 + 3b2)**, which were used for model building **(6a + b)**. The abundance of particles in the rotated and non-rotated state, as well as the occupancy of the A-site and tunnel exit are shown in a bar plot. FSC curves are plotted for both maps that were used for model building.

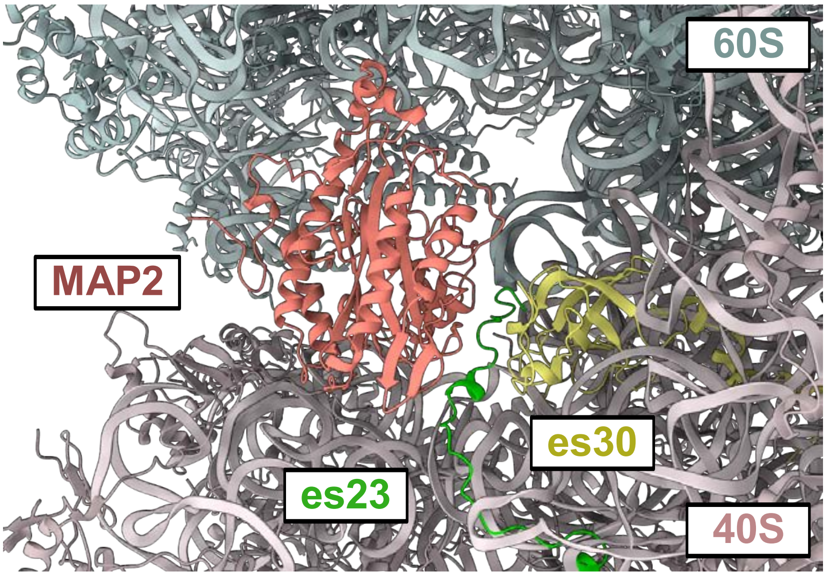

**Supplementary Figure 9:** *Ct*MAP2 binding in the A-site. MAP2 contacts 40S subunit protein es30 (yellow). In the presence of MAP2, the N-terminus of es23 (green) becomes ordered.

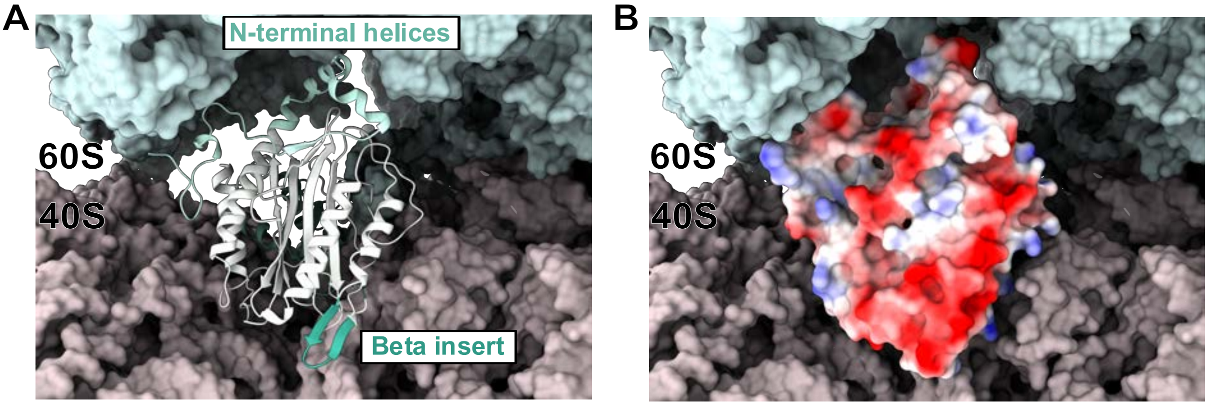

**Supplementary Figure 10:** Binding mode of *Ct*MAP2 in the ribosomal A-site. **(A)** Front view onto the A-site position of the ribosome. MAP2 binds in the interface between the 40S and 60S subunit. The N-terminal helices form strong interactions with the sarcin-ricin loop. The beta insert is not involved in binding. **(B)** The negatively charged beta insertion is solvent exposed and faces the entry of the ribosomal factor binding site.

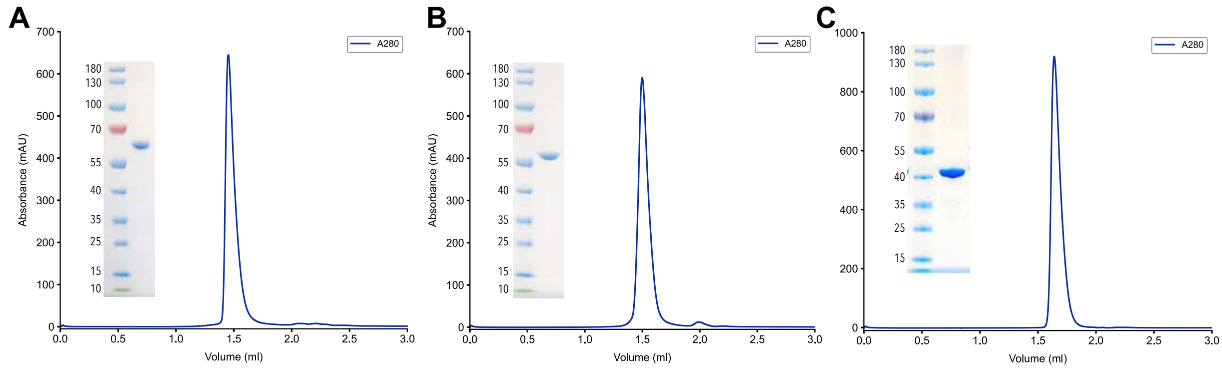

**Supplementary Figure 11:** Analytical SEC and SDS-gels of purified proteins used for cryo-EM sample preparation. SEC runs were performed on a S200/3.2-300 Increase column. **(A)** Purified HsMAP2 **(B)** *Ct*MAP2 and **(C)** *Ct*MAP2ΔN. Molecular weight of the protein marker is given in kDa.

### **Supplementary Table 1:** Cryo-EM data refinement statistics

| **Model** | ***hs*MAP2/80S** | ***ct*MAP2/80S** | ***ct*MAP2ΔN/80S** |
|---|---|---|---|
| *Data collection statistics* | | | |
| Microscope | Titan Krios | Titan Krios | Titan Krios |
| Camera | K2 | K3 | K2 |
| Voltage (kV) | 300 | 300 | 300 |
| Magnification | 84,000 | 105,000 | 84,000 |
| Total dose ($e^-$/Å$^2$) | 42 | 42.7 | 54.8 |
| Defocus range (μm) | -1.1 to -2.1 | -1.1 to -2.1 | -1.1 to -2.1 |
| Calibrated pixel size (Å) | 1.11 | 0.868 | 1.11 |
| *Refinement statistics* | | | |
| Refined particles | 215.768 | 61.734 | 119.160 |
| Resolution (Å) | 2.9 | 2.9 | 3.1 |
| Chains | 9 | 8 | 93 |
| Atoms | 12393 (Hydrogens: 0) | 14990 (Hydrogens: 0) | 208771 (Hydrogens: 0) |
| Residues | Protein: 960 Nucleotide: 214 | Protein: 991 Nucleotide: 329 | Protein: 12237 Nucleotide: 5230 |
| Water | - | - | - |
| Ligands | Co: 2 | - | ZNS: 3, B8N: 1, ZNL: 5 |
| Bonds (RMSD*) | | | |
| Length (Å) (# > 4σ) | 0.003 (0) | 0.004 (0) | 0.006 (7) |
| Angles (°) (# > 4 σ) | 0.638 (1) | 0.743 (13) | 0.614 (89) |
| MolProbity score | 1.34 | 1.62 | 1.68 |
| Clash score | 3.66 | 11.10 | 7.15 |
| Ramachandran plot (%) | | | |
| Outliers | 0.00 | 0.00 | 0.03 |
| Allowed | 3.06 | 2.25 | 4.17 |
| Favored | 96.94 | 97.75 | 95.80 |
| Rama-Z (Ramachandran Plot, Z-score, RMSD*) | | | |
| whole (N = 951) | 0.62 (0.25) | 0.33 (0.25) | -1.04 (0.07) |
| helix (N = 379) | 1.67 (0.26) | 1.67 (0.25) | -0.14 (0.08) |
| sheet (N = 107) | 1.60 (0.52) | 0.62 (0.43) | -0.15 (0.12) |
| loop (N = 465) | -0.98 (0.24) | -0.09 (0.28) | -1.15 (0.07) |
| Rotamer outliers (%) | 0.00 | 0.00 | 0.16 |
| Cβ outliers (%) | 0.00 | 0.00 | 0.00 |
| Peptide plane (%) | | | |
| Cis proline/general | 0.0/0.0 | 2.3/0.0 | 0.8/0.0 |
| Twisted proline/general | 0.0/0.0 | 0.0/0.0 | 0.2/0.0 |
| CaBLAM outliers (%) | 1.71 | 1.03 | 2.72 |
| ADP (B-factors) | | | |
| Iso/Aniso (#) | 12393/0 | 14990/0 | 208771/0 |
| min/max/mean | | | |
| Protein | 11.76/71.25/41.64 | 15.28/125.97/49.34 | 46.72/625.58/113.03 |
| Nucleotide | 18.95/124.23/44.59 | 24.96/235.84/10.57 | 47.08/689.64/103.76 |
| Ligand | 88.76/99.16/93.96 | - | 70.69/161.53/134.19 |
| Occupancy (%) | 100.00 | 100.00 | 100.00 |
| Model vs. Data (CC mask) | 0.84 | 0.77 | 0.87 |

* RMSD: root-mean-squared-deviation

**Supplementary Table 2:** X-ray data collection and refinement statistics for *Ct*MAP2ΔN

| | |
|---|---|
| *Data collection* | |
| Wavelength (Å) | 0.9737 |
| Resolution range (Å) | 46.53 - 1.3 (1.32-1.3) |
| Space group | P 21 21 21 |
| Unit cell (Å) | 59.94 73.81 82.05 90 90 90 |
| Total reflections | 1103571 (35025) |
| Unique reflections | 90066 (4362) |
| Multiplicity | 12.3 (8.0) |
| Completeness (%) | 99.9 (99.0) |
| Mean I/sigma (I) | 23.0 (2.1) |
| Wilson B-factor (Å$^2$) | 17.1 |
| $R_{pim}$ | 0.015 (0.288) |
| CC1/2 | 0.999 (0.879) |
| *Refinement* | |
| Resolution range (Å) | 40.47-1.3 (1.35-1.3) |
| Reflections used in refinement | 89942 (8840) |
| $R_{work}$ | 0.129 (0.198) |
| $R_{free}$ | 0.156 (0.234) |
| Number of non-hydrogen atoms | 3573 |
| macromolecules | 3043 |
| ligands | 20 |
| solvent | 522 |
| Protein residues | 371 |
| RMS (bonds) | 0.010 |
| RMS (angles) | 1.12 |
| Ramachandran favored (%) | 98.37 |
| Ramachandran allowed (%) | 1.63 |
| Ramachandran outliers (%) | 0.00 |
| Rotamer outliers (%) | 0.91 |
| Clashscore | 2.97 |
| Average B-factor (Å$^2$) | 24.43 |
| macromolecules | 22.39 |
| ligands | 36.16 |
| solvent | 36.15 |

Statistics for the highest-resolution shell are shown in parentheses.

$R_{free}$ is for 5% of all data.
